# Microglial cathepsin B promotes neuronal efferocytosis during brain development

**DOI:** 10.1101/2024.12.03.626596

**Authors:** Nicholas J Silva, Sarah Anderson, Supriya A Mula, Caroline C Escoubas, Haruna Nakajo, Anna V Molofsky

## Abstract

Half of all newborn neurons in the developing brain are removed via efferocytosis - the phagocytic clearance of apoptotic cells. Microglia are brain-resident professional phagocytes that play important roles in neural circuit development including as primary effectors of efferocytosis. While the mechanisms through which microglia recognize potential phagocytic cargo are widely studied, the lysosomal mechanisms that are necessary for efficient digestion are less well defined. Here we show that the lysosomal protease cathepsin B promotes microglial efferocytosis of neurons and restricts the accumulation of apoptotic cells during brain development. We show that cathepsin B is microglia-specific and enriched in brain regions where neuronal turnover is high in both zebrafish and mouse. Myeloid-specific cathepsin B knockdown in zebrafish led to dysmorphic microglia containing undigested dead cells, as well as an accumulation of dead cells in surrounding tissue. These effects where phenocopied in mice globally deficient for *Ctsb* using markers for apoptosis. We also observed behavioral impairments in both models. Live imaging studies in zebrafish revealed deficits in phagolysosomal fusion and acidification, and live imaging of cultured mouse microglia reveal delayed phagocytosis consistent with impairments in digestion and resolution of phagocytosis rather than initial uptake. These data reveal a novel role for microglial cathepsin B in mediating neuronal efferocytosis during typical brain development.

## Introduction

Microglia are the professional phagocytes of the brain and are essential participants in brain development.^1–3^ They survey the brain parenchyma in an activity-dependent manner and can rapidly phagocytose portions of cells or entire cells during development, plasticity, and in disease.^4,5^ Microglia are vastly outnumbered by neurons in the developing brain, yet they are extremely efficient phagocytes, eliminating nearly half of all cells produced in the cerebral cortex.^6^ The vast majority of this cell removal occurs via efferocytosis, a phagocytic mechanism for engulfing entire cells that have died by apoptosis.^7^ This is an energetically demanding process, particularly during development, when neuronal turnover is high. Therefore, defining the cellular mechanisms that promote efficient efferocytosis by microglia is highly relevant to understanding typical brain development as well as the pathogenesis many neurodevelopmental and neurodegenerative diseases.

Efferocytosis is a multistep process. Microglia first identify dead cell cargo via a series of “eat me” signals, including phosphatidylserine, which are present on the surface of apoptotic cells. ^7^ They then take up the cargo into a phagosome which then fuses with lysosomes to enable digestion within the phagolysosome.^7,8^ This fusion event generates an acidified phagolysosome where hydrolytic enzymes degrade proteins, nucleic acids, and other cellular material.^9^ The last stage of phagocytosis is resolution, during which the phagolysosome is compacted, its digestion products eliminated, and reusable elements including membranes recycled for subsequent rounds of phagocytosis.^10^ While the initial stages of microglial efferocytosis including recognition have been extensively studied, the mechanisms through which microglia efficiently digest and process phagocytic cargo are relatively less well understood. Some pathways have been identified, including vacuolar ATPases that promote acidification^11^, a glucose transporter that promotes compaction of the phagosome^12^, and Type I interferon signaling, which promotes whole cell phagocytosis through as yet undefined mechanisms.^13^ One clear outcome of these studies is that impairment at any stage of the digestion process can reduce the efficiency of phagocytosis, leading to a backup of undigested material.

Efficient digestion requires degradation of proteins in the phagolysosome. This process depends largely on cathepsins, the most abundant lysosomal proteases. There are three families of cathepsins, which include serine proteases (A and G), aspartic proteases (D and E) and cysteine proteases (B, C, F, H, L, O, S, V, X, and W).^14,15^ Cathepsin B is synthesized as a zymogen, an inactive enzyme that requires autocatalysis for full maturation within the lysosome. Under physiological states, cathepsin B is localized to the lysosome and endosome, where it aids in proteolysis of engulfed proteins.^14,15^ However, cathepsin B can be pathologic when released from the lysosome by physiological stress and in disease states.^16^ Moreover, cathepsin B is highly expressed in damage-associated microglia (DAMs) observed across disease settings, including in neurodegenerative diseases.^17–19^ We previously identified cathepsin B as highly expressed in developing zebrafish microglia and particularly enriched in microglia known to engulf apoptotic corpses in the optic tectum.^20^

Here, we studied the function of microglial cathepsin B in brain development using both zebrafish and mouse models. We found that cathepsin B expressing microglia were enriched in brain regions and time periods with high neuronal turnover including the zebrafish optic tectum and the murine somatosensory cortex. Zebrafish with myeloid-specific cathepsin B loss of function had microglia with a decreased ability to acidify phagosomes and deficient phagolysosomal fusion, together with an accumulation of dead cells in the brain and impaired locomotor behavior. Consistent with this, mice with loss of cathepsin B (*Ctsb^−/−^*) had impaired microglial phagocytosis of apoptotic cells. These mice also had an accumulation of apoptotic cells in layer 5 of the somatosensory cortex, a region with pronounced neuronal turnover at this age (postnatal day 5). Furthermore, cathepsin B deficiency in mice led to a decrease in cortical excitatory neuron density in the somatosensory cortex and tactile hypersensitivity. Taken together these data reveal a role for cathepsin B in promoting microglial efferocytosis during brain development and a requirement for cathepsin B in promoting neural circuit maturation.

## Results

### Cathepsin B is a lysosomal protease that is required for normal microglial numbers and morphology in developing zebrafish optic tectum

We previously identified two molecularly distinct populations of microglia in the developing zebrafish. One subset was enriched in a synapse-rich hindbrain region and expressed genes associated with microglial synaptic remodeling, whereas a microglial subset in the optic tectum (OT), a region with high neuronal turnover, was enriched for multiple cathepsins.^20^ The top differentially expressed gene enriched in OT microglia was the lysosomal protease cathepsin B (*ctsba*; **Figure 1A-B and S1A**). To define whether microglia had functional cathepsin B activity *in vivo*, we imaged live anesthetized fish after brain injection of the cathepsin B peptide substrate MR-(RR)2 (hereafter referred to as “Magic Red”), which becomes fluorescent after proteolytic cleavage by cathepsin B.^11^ We also delivered LysoTracker dye to label acidified compartments (**Figure 1C**).^11,21^ Microglia were identified by expression of a membrane localized GFP using the myeloid transgenic reporter *Tg(mpeg:GFP-CAAX)*. We observed cathepsin B activity (Magic Red+) only in microglia in the OT and found that cathepsin B activity was almost entirely observed within Lysotracker+ acidified compartments (**Figure 1D**).

**Figure 1:**
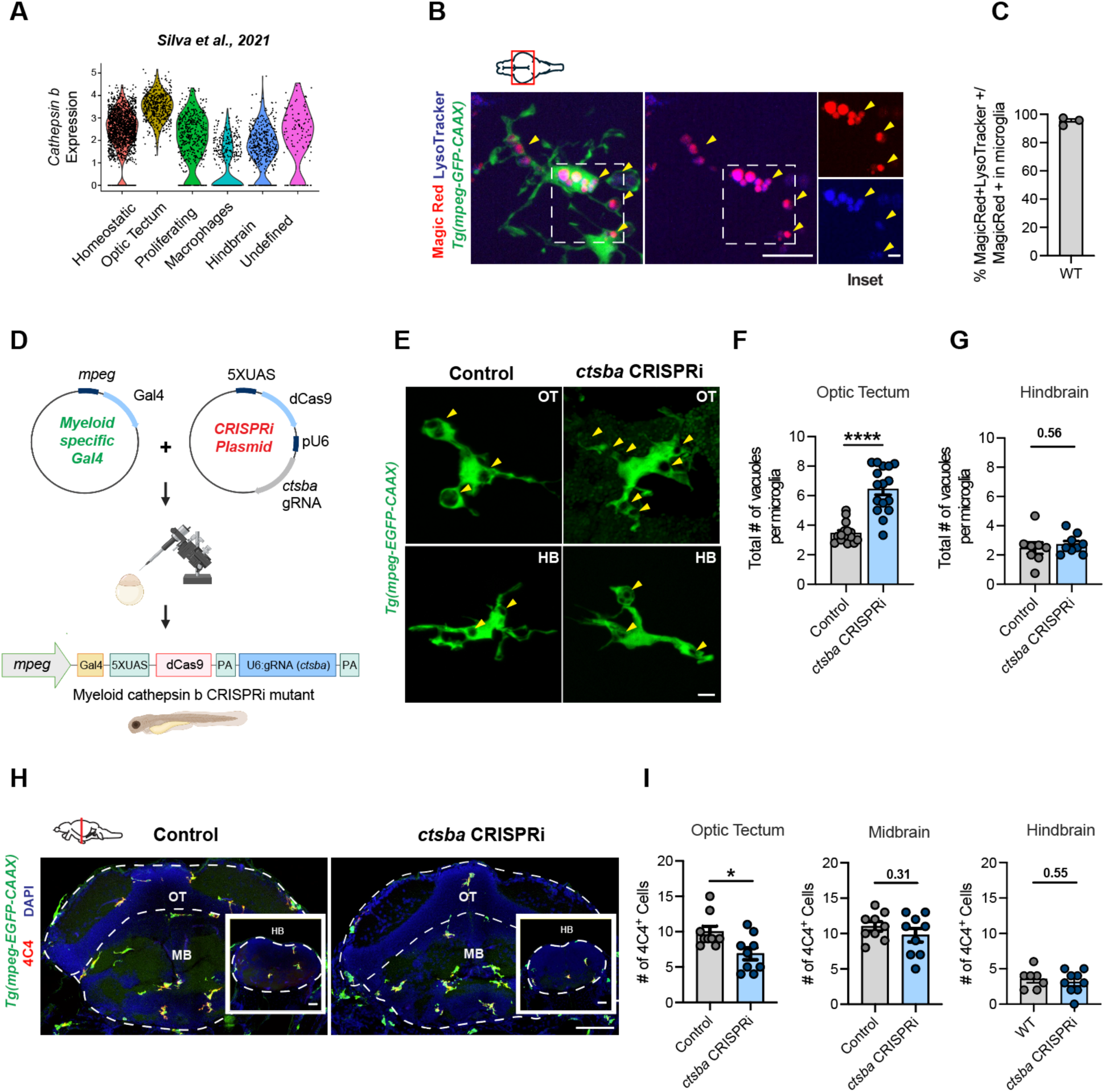
Cathepsin b is a lysosomal protease that is required for normal microglial numbers and morphology in developing zebrafish optic tectum. **(A)** Violin plot showing expression of the zebrafish gene for cathepsin B *(ctsba)* in microglial subsets identified by single cell RNA sequencing in *Silva et al., 2021*. Data from 28 days post fertilization (dpf). **(B)** Representative image of an OT microglia exposed to the cathepsin B substrate Magic Red-(RR)2 and the acidification marker LysoTracker at 10 dpf. Yellow arrowheads indicate cathepsin B+ lysosomes co-labeled with Magic Red and LysoTracker. Microglia defined by *Tg(mpeg1.1:GFP-CAAX)*. Scale bar = 10 µm. Inset scale bar = 5 µm **(C)** Quantification of the % cathepsin B-Magic Red signal colocalized with LysoTracker within GFP+ microglia. Dots represent means from 3 fish averaging 2-3 microglia per fish. **(D)** Strategy for myeloid specific CRISPRi knockdown of *ctsba*. Myeloid promoter, *mpeg1* was fused to Gal4-VP transactivator of upstream activation sequence (UAS). A kinase-dead Cas9 conjugated to kruppel associated box (KRAB) is expressed under a UAS promoter and a gRNA sequence is expressed under a ubiquitous U6 promoter for cell-specific gene silencing. **(E)** Representative images of control and *ctsba* CRISPRi mutants at 10 dpf. Arrowheads indicate phagosomal compartments. Scale bar = 10 µm. **(F)** Quantification of the number of vacuoles per microglia in OT control and *ctsba* CRISPRi mutants. Dots represent 16 fish/group, averaging 2-3 microglia per fish. Mann-Whitney U test, **** p <0.0001. **(G)** Quantification of the number of vacuoles per microglia in HB control and *ctsba* CRISPRi mutants. Dots represent 8 fish/group, averaging 2-3 microglia per fish. Welch’s t-test, ns (0.56). **(H)** Representative images of microglia co-labeled with *Tg(mpeg1.1:GFP-CAAX*) and 4C4 (microglia marker) in the OT and hindbrain (HB) from control and *ctsba* CRISRPi mutants at 10 dpf. Scale bar = 20 µm. **(I)** Quantifications of 4C4+ microglia in the optic tectum, midbrain, and hindbrain at 10 dpf. Dots = 9 fish/group averaging 2-3 microglia per fish. Welch’s t-test, * p <0.01 (OT), ns (MB), and ns (HB). See also Figure S1.

To study the function of cathepsin B during brain development we generated a cell-type specific knockdown in zebrafish microglia and myeloid cells using CRISPR interference (CRISPRi; **Figure 1E**). We expressed endonuclease-dead Cas9 protein (dCas9) and a guide RNA (gRNA) sequence targeting the *ctsba* gene under the ubiquitous U6 promoter and used the Gal4-UAS system^22^ to target expression to microglia and myeloid cells using the *mpeg* promoter. To validate knockdown of *ctsba,* we quantified Magic Red in controls (no transgene) and *ctsba* CRISPRi mutants. We observed a 3.45-fold reduction in Magic Red volume within microglia of *ctsba* CRISPRi mutants compared to controls (**Figure S1B-C**). We further quantified expression of *ctsba* from FACs isolated microglia by qPCR, which revealed a 37% reduction in *ctsba* transcript relative to housekeeping gene *ef1a* (**Figure S1D-E**). This assay demonstrates substantial though incomplete knockdown of cathepsin B in microglia.

We next examined the impact of cathepsin B knockdown on microglial morphology at 10 days post fertilization (dpf) in the OT. We observed that *ctsba* deficient microglia in the OT had more vacuoles compared to controls (defined as membrane enclosed compartments > 2 µm in diameter; **Figure 1E-F**). There was no difference in microglial volume (**Figure S1F**). Interestingly, vacuole number and microglia volume were unchanged in synapse-associated hindbrain microglia (**Figure 1G and S1G**). We also observed a ~25% reduction in the number of microglia in the OT but no differences in the midbrain and hindbrain as assessed by the microglia-specific marker 4C4 (Galectin 3)^23^ (**Figure 1H-I**). Together, our results suggest that cathepsin B is required for typical microglial morphology and numbers in the OT, where neuronal turnover is high, but is dispensable in synapse rich midbrain and hindbrain regions.

### Microglial Cathepsin B restricts the accumulation of dead cells and impacts locomotor behavior

Microglia in the OT are critical for efferocytosis of neurons that die during the process of neurogenesis and neuronal turnover.^24–26^ To examine if microglial cathepsin B was necessary for this process, we performed TUNEL staining (terminal deoxynucleotidyl transferase-mediated deoxyuridine triphosphate nick end labeling) to detect dead cells in control and microglial *ctsba* CRISPRi mutants. We found that *ctsba* CRISPRi mutants had an increase in the total number of TUNEL+ cells in the OT (**Figure 2A-B**). We also observed an accumulation TUNEL+ cells outside and within individual microglia in the OT (**Figure 2C-F**). We obtained concordant results using the cathepsin B cell-permeable inhibitor, CA-074Me^27^ that was delivered at a dose of 100 µm by swimming the fish for 3 days with daily replacements of the drug (**Figure S2A-C**). Together these results suggest that microglial cathepsin B promotes clearance of dead cells in the developing brain.

**Figure 2:**
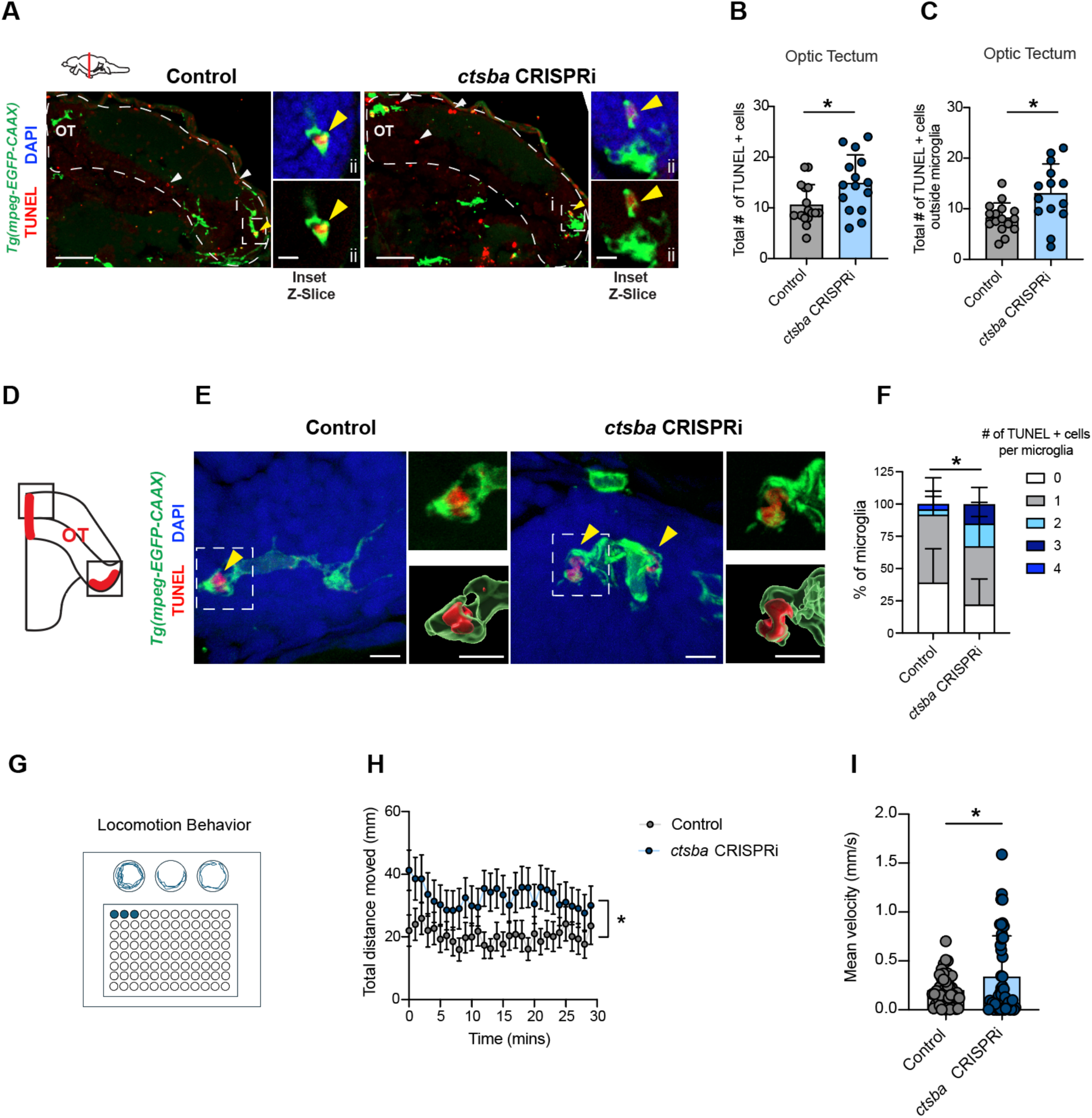
Microglial Cathepsin B restricts the accumulation of dead cells and impacts locomotor behavior. **(A)** Representative images of microglia labeled with *Tg(mpeg:GFP-CAAX)* and apoptotic debris detected by TUNEL (terminal deoxynucleotidyl transferase-mediated deoxyuridine triphosphate nick end labeling) from controls and *ctsba* CRISPRi mutants in the OT. White arrowheads indicate TUNEL+ cells outside of microglia. Yellow arrowheads in the inset indicate TUNEL+ cells in microglia. (i) Scale bar = 50 µm. (ii) Inset scale bar = 5 µm. **(B)** Quantification of total TUNEL positive cells in the optic tectum at 10 dpf. Dots = 15 fish/group averaging 2-3 microglia per fish. Welch’s T test, *p < 0.0228. **(C)** Quantification of percent TUNEL positive cells outside microglia from controls and *ctsba* CRISPRi mutants in the OT. Dots = 15 fish/group averaging 2-3 microglia per fish. Welch’s T test, *p < 0.0126. **(D)** Schematic diagram of regions examined in E-F. Red shading indicates regions of neuronal turnover in the OT. **(E)** Representative images of microglia containing TUNEL+ debris (yellow arrowheads_ from controls and *ctsba* CRISPRi mutants. Dotted line indicates inset region. Insets show raw image (top) and 3D reconstruction (bottom). Scale bar = 5 µm. Inset scale bar = 8 µm. **(F)** Quantification of the number of TUNEL positive cells within microglia. Data represented as stacked bar graphs. Data calculated from means of 12 fish/group averaging 2-3 microglia per fish. Fisher’s exact test, *p < 0.0263. **(G)** Schematic diagram of 96 well plate used to swim larvae and record locomotion. **(H)** Locomotion behavior quantified as distance traveled (mm) from control and *ctsba* CRISPRi mutants. Dots = mean value per fish from 44 fish/group. 2-way RM ANOVA, * p < 0.05, F (1,86) = 4.041. **(I)** Mean velocity in control and *ctsba* CRISPRi mutants. Welch’s T test, 0.0421. Dots = mean value per fish from 44 fish/group. See also Figure S2.

To assess whether these defects in microglial phagocytic function and dead cell clearance were associated with broader impacts on brain function, we observed spontaneous locomotion of zebrafish larvae using a high throughput automated imaging platform (**Figure 2G**).^28^ Following habituation to the recording platform, we recorded spontaneous movement for 30 mins. We found that *ctsba* CRISPRi mutants were hyperactive compared to controls as evidenced by an increase in total distance traveled and increased mean velocity (**Figure 2H-I**). Thus, myeloid-specific depletion of cathepsin B leads to altered locomotor behavior.

### Cathepsin B promotes phagolysosomal fusion and phagosome acidification

Efferocytosis, or engulfment of entire apoptotic cells, requires a robust and effective process of phagocytosis and digestion. An essential step in this process is fusion of the phagosome containing phagocytosed material with an acidic lysosome filled with pH sensitive proteases like cathepsins.^8,9^ The site of this phagolysosome fusion is marked by clustering of Rab7 protein, a small GTPase that promotes multiple steps of the phagocytic process (**Figure 3A**).^29^ To examine how Cathepsin B signaling regulates microglial phagocytosis, we quantified acidified compartments using LysoTracker (**Figure 3B**).^11,21^ We observed that despite the marked increase of phagocytic vacuoles in cathepsin B deficient microglia (**Fig. 1F**), there was a significant reduction in the percentage that were acidified (**Figure 3C**) with a trend towards fewer total acidified compartments (**Figure S3A**). This did not appear to be driven by a reduction in lysosomes, as the number of putative lysosomes did not differ between wildtype and *ctsba* CRISPRi mutants (defined as LysoTracker+ and ≤ 2 µm in diameter; **Figure S3B**).

**Figure 3:**
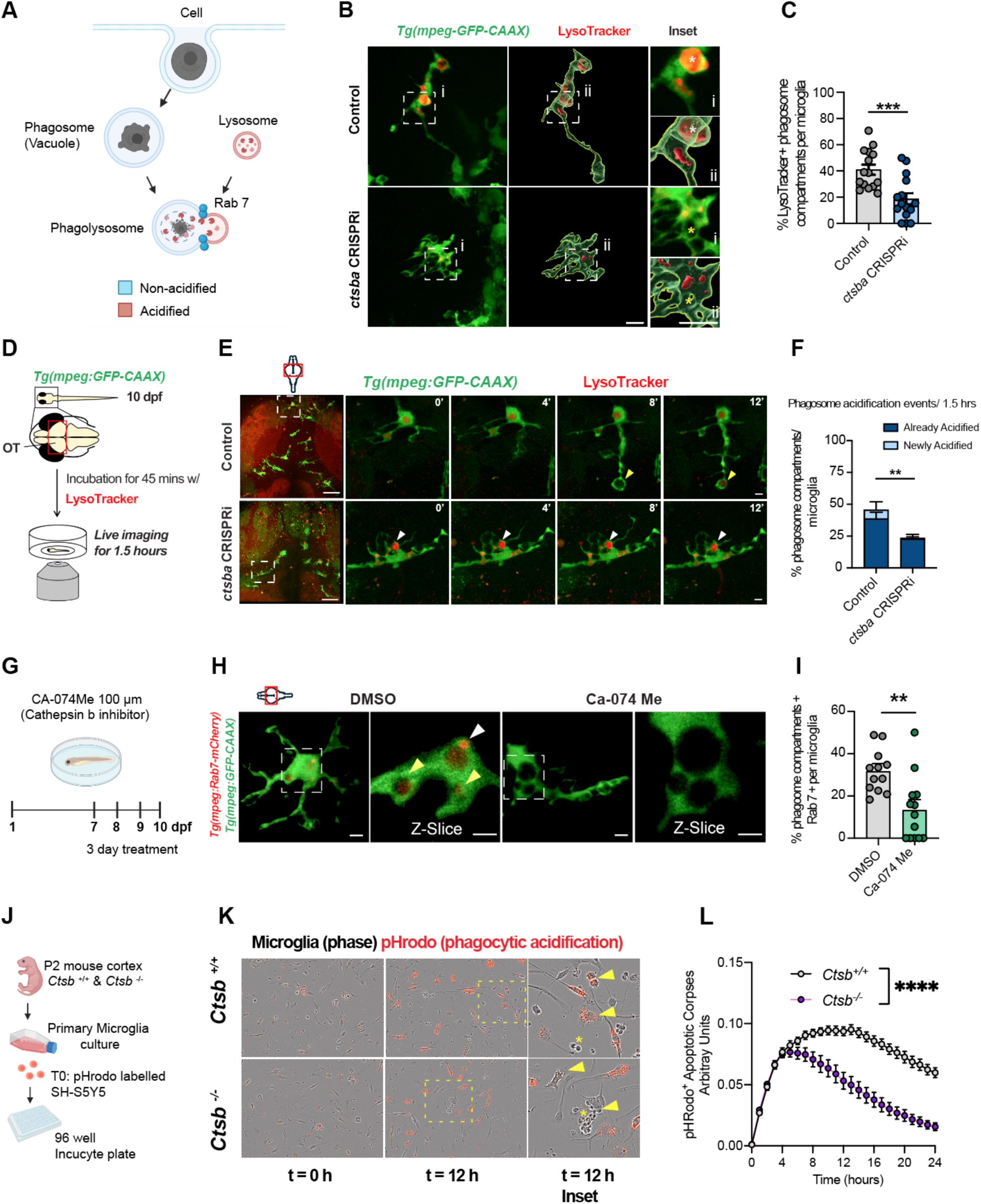
Cathepsin B promotes phagolysosomal fusion and phagosome acidification. **(A)** Schematic of stages of phagocytosis including phagosome formation and Rab 7-dependent phagolysosomal fusion. **(B)** Representative images of microglia Tg(*mpeg1.1-GFP-CAAX)* and LysoTracker signal in control and *ctsba* CRISPRi mutants at 10 dpf. Scale bar *=* 7 µm. Inset: White asterisk indicates LysoTracker + phagosome compartment and yellow asterisk indicates LysoTracker – phagosome compartment. Scale bar *=* 7 µm. **(C)** Quantification of the percent acidified phagosome compartments per microglia in control and *ctsba* CRISPRi mutants at 10 dpf. Dots represent16 fish/group averaging 2-3 microglia per fish. Welch’s t-test, *** p <0.0004. **(D)** Schematic of zebrafish soaked in LysoTracker and live-imaging experiment. **(E)** Time series of microglia (green, *Tg(mpeg1.1:EGFP-CAAX*)) and acidified compartments (Red, LysoTracker) in the zebrafish optic tectum at 10 dpf. Low power view (far left) includes outlined box which features representative microglia that was tracked over time in the subsequent insets. Yellow arrowhead indicates a newly acidified phagosome, white arrowhead is an already acidified phagosome. Scale bar = 20 µm. **(F)** Distribution of acidification events during the 90-minute imaging session for control and *ctsba* CRISPRi mutants at 10 dpf. Data calculated from means of 6-5 fish/group averaging 8-10 microglia per fish. Fisher’s exact test, ** p < 0.0065. **(G)** Schematic diagram of the pharmacological treatment with CA-074Me (Cathepsin B inhibitor, 100 µm) vs. DMSO control delivered over 3 days to fish water from 7-10 dpf. **(H)** Representative images of microglia (green, *Tg(mpeg1.1:EGFP-CAAX)* and Rab 7 signal (red, *Tg(mpeg:Rab7-mcherry))* at 10 dpf. White arrowhead indicates Rab7-mCherry protein. Yellow arrowheads indicate phagosomes that are Rab7-mCherry positive. Scale bar = 10 µm. **(I)** Quantification of the percentage of phagosomes that are positive for Rab 7-mCherry expression within microglia in DMSO and CA-074Me Dots = 10 fish/group averaging 2-3 microglia per fish. Welch’s T-test, ** p <0.0018. **(J)** Schematic of *in vitro* assay to quantify the impact of cathepsin B function on microglial phagocytosis of pHrodo-labeled apoptotic SH-S5Y5 cells by multiwell live cell imaging. SH-S5Y5 cells, added simultaneously to both groups at time = 0 (T0). **(K)** Representative images of primary microglial cultures at time = 0 and 12 from *Ctsb ^+/+^* and *Ctsb ^−/−^* groups. Yellow dashed box corresponds to insets. Yellow arrows show acidified SH-S5Y5 cells within microglia. Yellow asterisk shows undigested SH-S5Y5 cells. Scale bar = 10 µm. **(L)** Representative intensity curves from *Ctsb ^+/+^* and *Ctsb ^−/−^* primary microglial cultures fed with pHrodo ^+^ labeled apoptotic cells at T=0-24 h, light grey (*Ctsb ^+/+^*) and (*Ctsb ^−/−^*). 2-way RM ANOVA, **** p < 0.0001, F (1, 24) = 36.04). See also Figure S3.

We next examined the dynamics of phagolysosomal fusion by using live imaging to quantify real-time acidification events within microglia (**Figure 3D-E**). We classified microglial phagocytic vacuoles over the 90-minute imaging period as ‘Already acidified’ or ‘Newly acidified’ based on whether they were or became LysoTracker+ during the imaging window (**Videos S3A-B**). We observed that *ctsba* CRISPRi mutant microglia had fewer already acidified and newly acidified compartments (**Figure 3F**). The morphology of these events also differed. Control microglia extended long processes that formed phagocytic compartments that subsequently acidified. In contrast, *ctsba* CRISPRi mutant microglia had minimal process motility, and most acidified compartments were observed near the microglial soma. These data suggest that cathepsin B promotes efficient phagosome acidification.

To directly test whether these defects in phagosome acidification reflected impaired phagolysosomal fusion, we quantified Rab 7, which clusters at fusion sites (**Figure 3A**). using the transgenic reporter *Tg(mpeg:Rab7-mCherry)*^21^. We used the inhibitor CA-074Me to globally suppress cathepsin B function (**Figure 3G**). We observed that cathepsin B blockade significantly decreased both the percent of microglia containing Rab7-mCherry+ phagosomes and the total Rab7-mCherry volume in microglia (**Figure 3H-I and S3C**). These results indicate that the acidification defects following *ctsba* inhibition may be in part due to impaired fusion between the phagosome and the lysosome.

To further examine the role of cathepsin B in engulfment and acidification using an orthogonal approach, we cultured primary microglia from *Ctsb^−/−^* deficient mice and their littermate controls (*Ctsb^+/+^*) and used an *in vitro* phagocytosis assay to track and quantify microglial digestion of apoptotic corpses labeled with pHrodo, a dye which fluoresces at acidic pH (**Figure 3J**).^13^ We found that *Ctsb^−/−^* microglia initially engulfed apoptotic corpses at a similar rate to controls (**Figure 3K**). However, they achieved their peak engulfment capacity significantly sooner than control cells (at ~5 hours vs. 9 hours) and engulfed significantly less than their wild type counterparts over the 24-hour period. (**Figure 3M and S3D**). These data suggest that cathepsin B is dispensable for the initial stages of efferocytosis, but becomes progressively more necessary at stages when digestion and resolution of the phagolysosome are required to enable subsequent rounds of digestion.

### Cathepsin B promotes microglial efferocytosis of neurons during mouse brain development

Cathepsin B is a core microglia gene expressed across species and highly enriched in microglia relative to other brain cells in both human and mouse (**Figure S4A-B**).^30,31^ We therefore examined whether the developmental function of cathepsin B was conserved in mice. We focused our studies on the early postnatal rodent barrel cortex (postnatal day 5, P5), where we previously observed microglial-mediated neuronal elimination selectively within cortical layer 5 (L5).^13^ Consistent with our findings in fish, we found that cathepsin B protein was primarily detected in microglia, using an antibody validated against *Ctsb^−/−^*tissue (**Figure S4C**). We also observed that cathepsin B was strongly enriched in L5 relative to other cortical layers, consistent with a role in cellular engulfment (**Figure 4A-B**). Within L5 microglia, we found that cathepsin B staining correlated with CD68, a membrane protein found on lysosomes, late endosomes, and phagolysosomes (**Figure 4C**).^32^ We then examined mice globally deficient for cathepsin B (*Ctsb^−/−^*). We found that *Ctsb^−/−^*had more CD68+ compartments per microglia compared to *Ctsb^+/+^*littermate controls (**Figure 4D-E**), as well as an increase in CD68 volume (**Figure S4D**). Microglia number was unchanged (**Figure S4E-F**). These data indicate that cathepsin B promotes microglial phagocytic function in mice.

**Figure 4:**
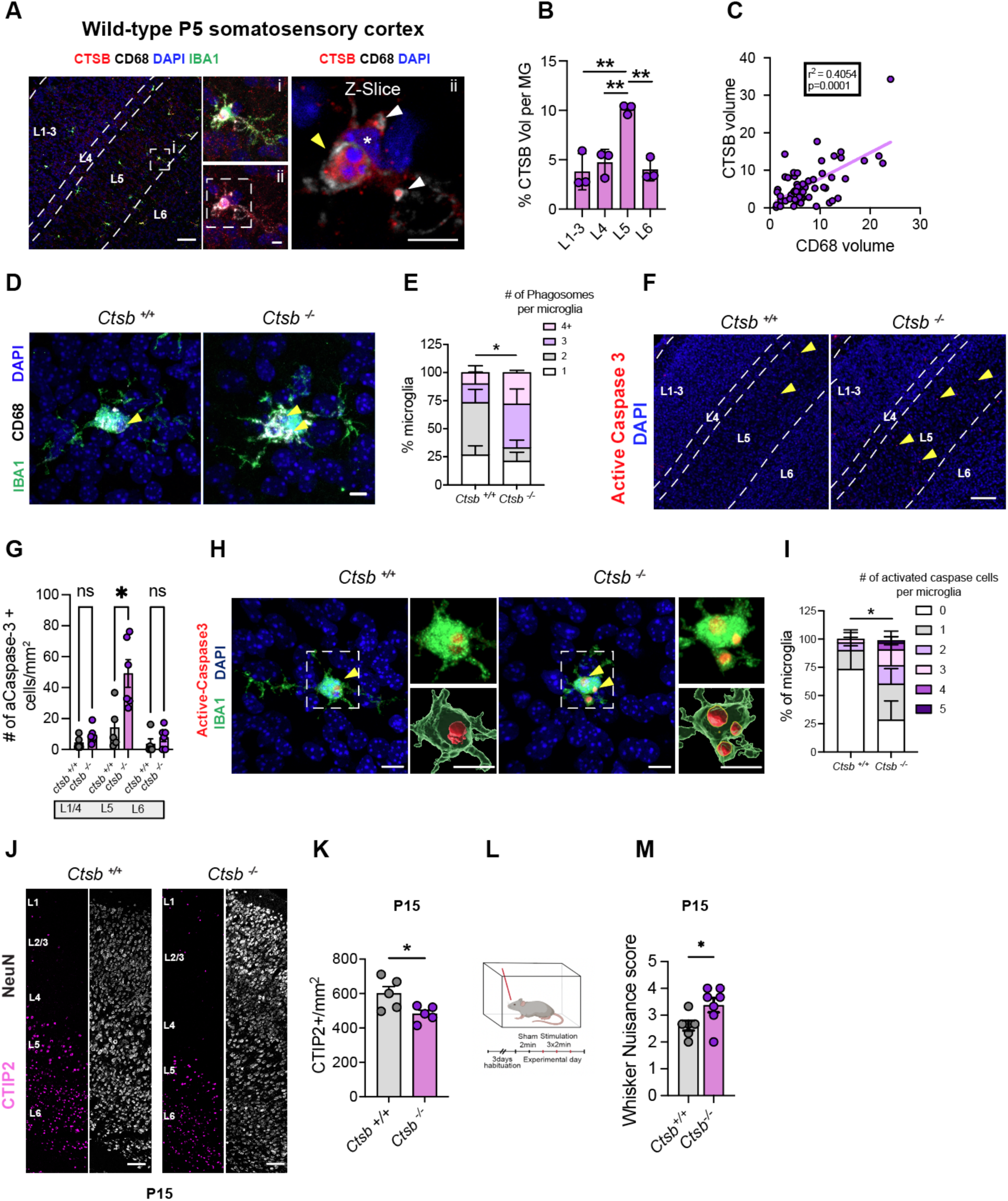
CTSB promotes microglial efferocytosis during mouse brain development. **(A)** Representative images from the somatosensory cortex in postnatal day 5 (P5) mice, showing antibody staining for microglia (IBA1+) cathepsin B, CD68+ phagocytic compartments, and DAPI nuclear stain. Dashed lines delineate cortical layers. Scale bar = 100 µm. (i) Inset indicates a single microglia in layer 5 (L5) labeled for all four colors (top), and (ii) without IBA1 (bottom). Scale bar = 5µm. (ii) inset shows a z-slice through the microglia highlighting cathepsin B in CD68+ phagolysosomal compartments. Yellow arrowhead shows a phagolysosome, white arrowheads indicate lysosomes, and asterisk is the microglia nucleus. Scale bar = 5 µm. **(B)** Quantification of the percentage of cathepsin B volume per microglia in cortical layers 1-6 at P5. One way ANOVA with Tukey’s multiple comparisons, **p<0.01. **(C)** Correlation between CD68 volume and Cathepsin B volume per microglia at P5 (n=59 cells from 3 mice, r Spearman correlation coefficient = 0.40, p<0.0001). **(D)** Representative images of CD68+ phagolysosomal compartments co-stained with microglia (IBA1+) from *Ctsb ^+/+^* and *Ctsb ^−/−^*mice at P5. **(E)** Quantification of the number of phagosomes per microglia. Phagosomes were defined as CD68+ vacuoles ≥3 µm in diameter. Data calculated from means of 5-4 mice/group averaging 2-3 microglia per mouse. Fisher’s exact test, * p <0.0462. **(F)** Representative images of active caspase 3 staining in the somatosensory cortex of layers L1-L6 from *Ctsb ^+/+^* and *Ctsb ^−/−^* mice at P5. (n= 5 mice for *Ctsb ^+/+^* and n= 6 mice for *Ctsb ^−/−^*). **(G)** Quantification of active-caspase 3 positive cells within layers L1-L6 of the barrel cortex at P5. (n= 5 mice/group). 2-way RM ANOVA with Sidak’s multiple comparisons, p* < 0.0349. **(H)** Representative images of active-caspase 3+ cells within microglia in L5 from *Ctsb ^+/+^* and *Ctsb ^−/−^* mice. Yellow arrowheads indicate active-caspase 3 positive cells inside microglia. Inset shows raw image (top) and 3D reconstruction (bottom). Scale bar = 20 µm. **(I)** Quantification of the number of active-caspase 3+ cells within microglia from *Ctsb ^+/+^* and *Ctsb ^−/−^* mice. Data calculated from 4-5 mice/group averaging 2-3 microglia per mouse. Fisher’s exact, **p < 0.0066. **(J)** Representative images of CTIP2 and NeuN positive neurons in the somatosensory cortex of P15 *Ctsb ^+/+^* and *Ctsb ^−/−^* mice. Scale bar = 50 µm. **(K)** Total CTIP2^+^ neuron density per mm^2^ across all cortical layers in *Ctsb ^+/+^* and *Ctsb ^−/−^* mice, P15 (n=5 mice per group). Welch’s t-test, *p=0.0436. **(L)** Schematic of whisker nuisance assay. A higher score reflects increased aversive response to tactile stimulus (see methods). **(M)** Whisker nuisance score in P15 *Ctsb ^+/+^* and *Ctsb ^−/−^* mice (n= 5 *Ctsb ^+/+^* mice and n= 8 *Ctsb ^−/−^*mice). Welch’s t-test, *p < 0.0375. See also Figure S4.

To determine whether Cathepsin B was required for elimination of apoptotic cells in the murine cortex, we quantified apoptosis using an antibody for active caspase-3.^33^ We found that *Ctsb^−/−^*mutants had an increase in total apoptotic cells, which was specific to L5 (**Figure 4F-G**). We also observed an accumulation of apoptotic material within microglia (**Figure 4H-I**), as well as an increase in TUNEL+ material (**Figure S4G-H**). Taken together, our results indicate that cathepsin B promotes microglial efferocytosis of neurons in the developing mouse brain.

Finally, we examined the impact of cathepsin B on mouse cortical development and function. Since our effects were predominantly in L5, we quantified the excitatory neuron subtypes in this region using the markers CTIP2 and SATB2.^13^ During peak apoptosis (P5) we did not observe a difference in the number of CTIP2+, SATB2+, or double positive neurons (**Figure S4I-L**). However, by P15 the density of CTIP2^+^ neurons in L5/6 were significantly reduced (**Figure 4J-K**). There was no change in the overall number of total neurons (NeuN^+^) or upper layer excitatory neurons (SATB2+) in *Ctsb ^−/−^* mice (**Figure 4J and S4M**), suggesting this effect was specific to layer 5 excitatory neurons, where apoptotic cell death and phagocytic microglia were observed at P5. To test if cathepsin B impacted somatosensory behavior, we used a whisker nuisance assay at P15 to measure aversive behavioral responses to tactile stimulation using a standardized rating scale^34,35^, as we previously optimized in juvenile mice.^13^ We observed a significant increase in tactile hypersensitivity in *Ctsb ^−/−^* mice (**Figure 4L-M**). Taken together, these data reveal a role for cathepsin B-mediated microglial efferocytosis in regulating neuronal numbers and somatosensory circuit function during mouse cortical development.

## Discussion

Microglial efferocytosis is essential to brain development and regeneration.^7^ Our study identifies microglial cathepsin B as a novel regulator of microglial efferocytosis that is necessary to eliminate neurons during brain development.

A major finding from our study is that cathepsin B deficiency leads to a buildup of non-acidified vacuoles within microglia and a reduction in phagolysosomal fusion and acidification. This raises the question of how a protease that is largely active at acidic pH prevents a buildup of non-acidified compartments. One explanation is that impaired proteolysis leads to a ‘traffic jam’ in the phagocytic process by impairing phagosome resolution. Resolution of the phagolysosome is a critical step in phagocytosis. It occurs following digestion of phagolysosomal contents via a series of fragmentation events that generate smaller phagosome-derived vesicles.^36^ Resolution is vital to regenerating resources needed for subsequent rounds of phagocytosis, including regeneration of the lysosome pool.^10^ Resolution requires efficient cargo degradation^10,37^, therefore loss of cathepsin B would be predicted to impair resolution. Consistent with this hypothesis, our in vitro studies showed that the initial stages of phagocytosis proceeded normally, and impaired apoptotic cell uptake only became evident four hours after addition of phagocytic cargo (**Figure. 3K-L**). This places cathepsin B in a similar category to other mechanisms that promote orderly and efficient phagocytosis, including v-ATPases that promote acidification^11^, mechanisms of gastrosome maturation^12^, and cation channels that are required for phagosome resolution.^10,38^

Another key finding from our study is that loss of microglial cathepsin B led to a buildup of dead cells in developing brain that was closely correlated with the accumulation of dysfunctional microglia. Our mouse studies clearly identified these cells as apoptotic based on active caspase-3 staining (**Figure. 4**). These two interrelated phenotypes in microglia and dead cells after cathepsin B knockdown are consistent with other studies that have studied pathways involved in lysosomal function.^21,39,40^ For example, microglia lacking the lysosomal GTPase RagA have an increase in phagosomes and an accumulation of neuronal debris.^21,39^ Mutations of the *mcoln1* in zebrafish and drosophila lead to increased neuronal death ^41,42^ and visual and motor deficits also seen in patients with mucolipidosis type IV.^43^ These findings suggest that cathepsin B’s role in microglia phagolysosome function is necessary for healthy brain development, consistent with our findings of behavioral deficits in both zebrafish and mouse models of cathepsin B deficiency.

Finally, it is notable that the requirement for cathepsin B in microglial efferocytosis was most pronounced in regions with high neuronal turnover, including the zebrafish optic tectum (relative to midbrain and hindbrain) and the mouse somatosensory cortex in layer 5 (relative to other cortical layers). These data suggest that while cathepsin B is not the only mechanism for mediating microglial phagocytosis, it is particularly relevant in settings of high phagocytic demand, such as whole cell efferocytosis. This is consistent with other studies. Mice lacking cathepsins B and L show an increase in apoptosis and a decrease of neurons in some brain regions ^44^ and *Ctsb ^−/−^* mouse embryonic fibroblast exhibit NPC disease-like cholesterol accumulation and lysosomal dysfunction.^45^ And while cathepsin B itself is not associated with any known human lysosomal storage disorder, Cathepsin B activity is increased in Niemann-Pick type C disease^46,47^, suggesting that it could play some compensatory role. This implies that unlike in neurodegeneration, where cathepsin B has been proposed to be pathogenic, ^48^ it is possible that promoting cathepsin B function during development could be therapeutic in neurodevelopmental diseases.

## Limitations of the study

While cathepsin B is highly specific to microglia within the brain, our knockdown of cathepsin B in zebrafish also targets other myeloid cells, thus we cannot rule out that global phenotypes, such as behavior, are partly due to peripheral macrophage effects. Similarly, our mouse mutant targets cathepsin B in all cells including neurons.

## Acknowledgements

We are grateful to members of the Molofsky Lab for helpful comments on the manuscript, to Drs. Will Talbot and Harini Iyer for their expert advice, and to Louie Ramos, Stephanie Gilbert, and William Figueroa for the support of the CVRI Zebrafish Core facility. Thanks to Dr. Will Talbot for the Tg(*mpeg1.1:Rab7-mCherry*) transgenic reporter fish^21^, Dr. Jason Cyster for sharing *Ctsb^−/−^* mice that was made from Deussing J et al^49^ study, Dr. Cody smith for the CRISPRi backbone plasmid and Dr. Sarah Kucenas for the *mpeg1.1*:Gal-VP tol2 plasmid, and Dr. Kari Herrington at the UCSF Center for Advanced Light Microscopy (CALM) for technical imaging contributions.

## Author contributions

Conceptualization: N.J.S and A.V.M; Methodology, N.J.S., S.M., S.A., and C.C.E; Investigation, N.J.S., S.M., S.A., and C.C.E; Writing – Original Draft, N.J.S; Writing – Review & Editing, all co-authors; Funding Acquisition, A.V.M. and N.J.S. Resources, A.V.M and N.J.S.; Supervision, A.V.M. and N.J.S

## Data and materials availability

Supplement contains additional data. All data needed to evaluate the conclusions in the paper are present in the paper or the Supplementary Materials.

---

**Figure S1:**
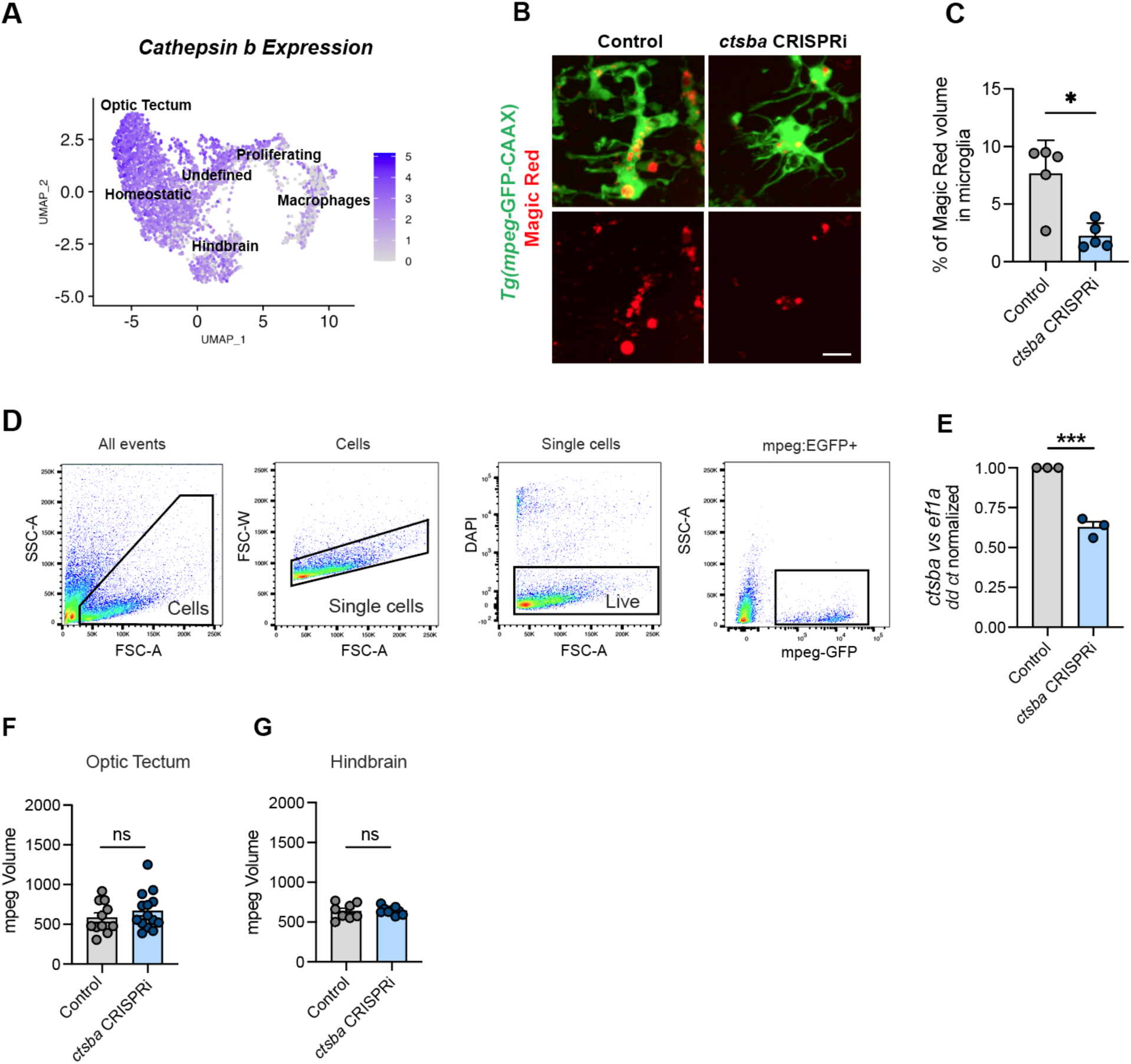
Validation of *ctsba* CRISPRi mutants, related to Figure 1. **(A)** Feature plot cathepsin B gene (*ctsba*) expression in UMAP space from 28 dpf zebrafish microglia, from *Silva et al, 2021*. **(B)** Representative images of cathepsin B-activity based on cathepsin B peptide substrate Magic Red-(RR)2 in microglia from control and *ctsba* CRISPRi mutants. Scale = 10 µm. **(C)** Quantification of %Magic Red within microglia from wild-type and *ctsba* CRISPRi mutants. Dots = 5 fish/group averaging 2-3 microglia per fish. Welch’s t-test, * p < 0.010. **(D)** Flow cytometry gating strategy to isolate *mpeg1.1*-EGFP+ myeloid cells for qPCR from control and *ctsba* CRISPRi mutants using whole larvae at 10 dpf. **(E)** qPCR for *ctsba* expression from flow sorted *mpeg+* cells from control and *ctsba* CRISPRi mutant whole larvae at 10 dpf. Expression levels are normalized to housekeeping gene *ef1a*. (Dots represent independent replicates, each pooled from 10-15 fish). Welch’s t-test, *** p < 0.0005. **(F)** Quantification of OT mpeg volume from control and *ctsba* CRISPRi mutant microglia. Dots represent 12 fish/group averaging 2-3 microglia per fish. ns, p=0.0574. **(G)** Quantification of HB mpeg volume from control and *ctsba* CRISPRi mutant microglia. Dots represent 9 fish/group averaging 2-3 microglia per fish. ns, p=0.8546.

**Figure S2:**
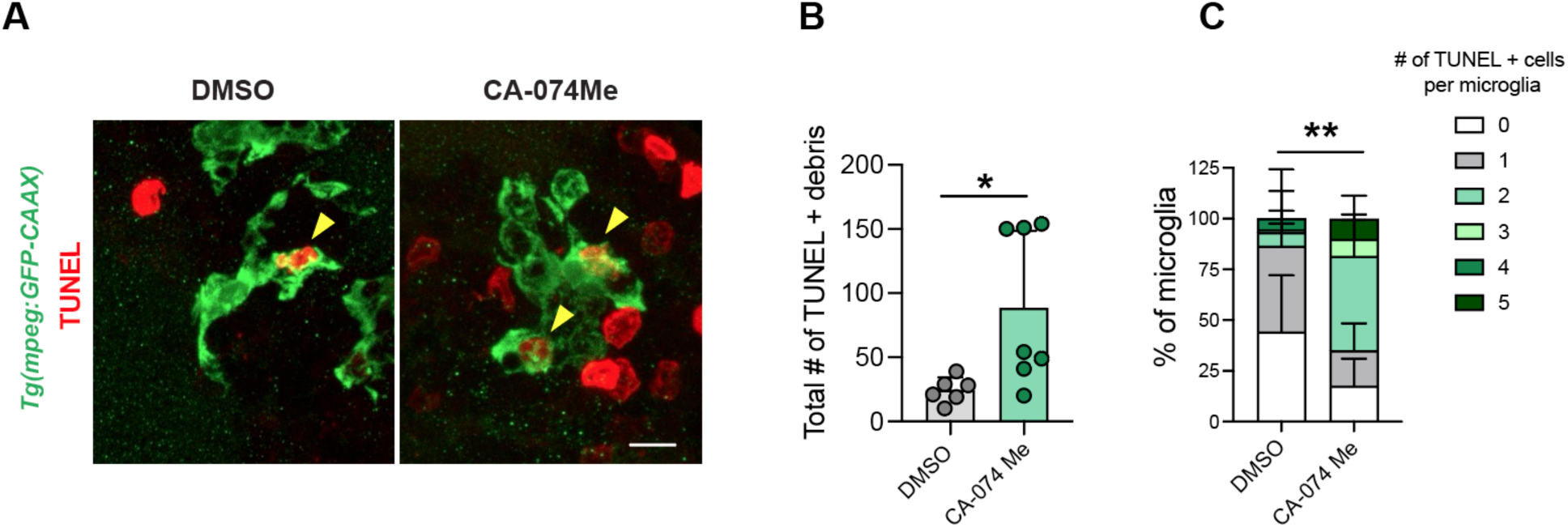
Impact of cathepsin B pharmacological inhibition on cell death and microglial efferocytosis, related to Figure 2. **(A)** Representative images of dead cells detected by TUNEL (terminal deoxynucleotidyl transferase-mediated deoxyuridine triphosphate nick end labeling) and microglia *Tg(mpeg:EGFP-CAAX*) from DMSO and CA-074Me treated groups. Yellow arrowheads indicate TUNEL+ cells within microglia. Scale bar =10 µm. **(B)** Quantification of the total number of TUNEL+ cells in OT from DMSO and the cathepsin B inhibitor CA-074Me treated groups at 10 dpf. Dots = 6 fish/group, averaging 2-3 microglia per fish. Welch’s t-test, * p < 0.0301. **(C)** Quantification of the number of TUNEL+ cells within microglia from DMSO and CA-074Me treated groups in the optic tectum at 10 dpf. Data calculated from means of 6 fish/group averaging 2-3 microglia per fish. Fisher’s exact test, ** p < 0.0052.

**Figure S3:**
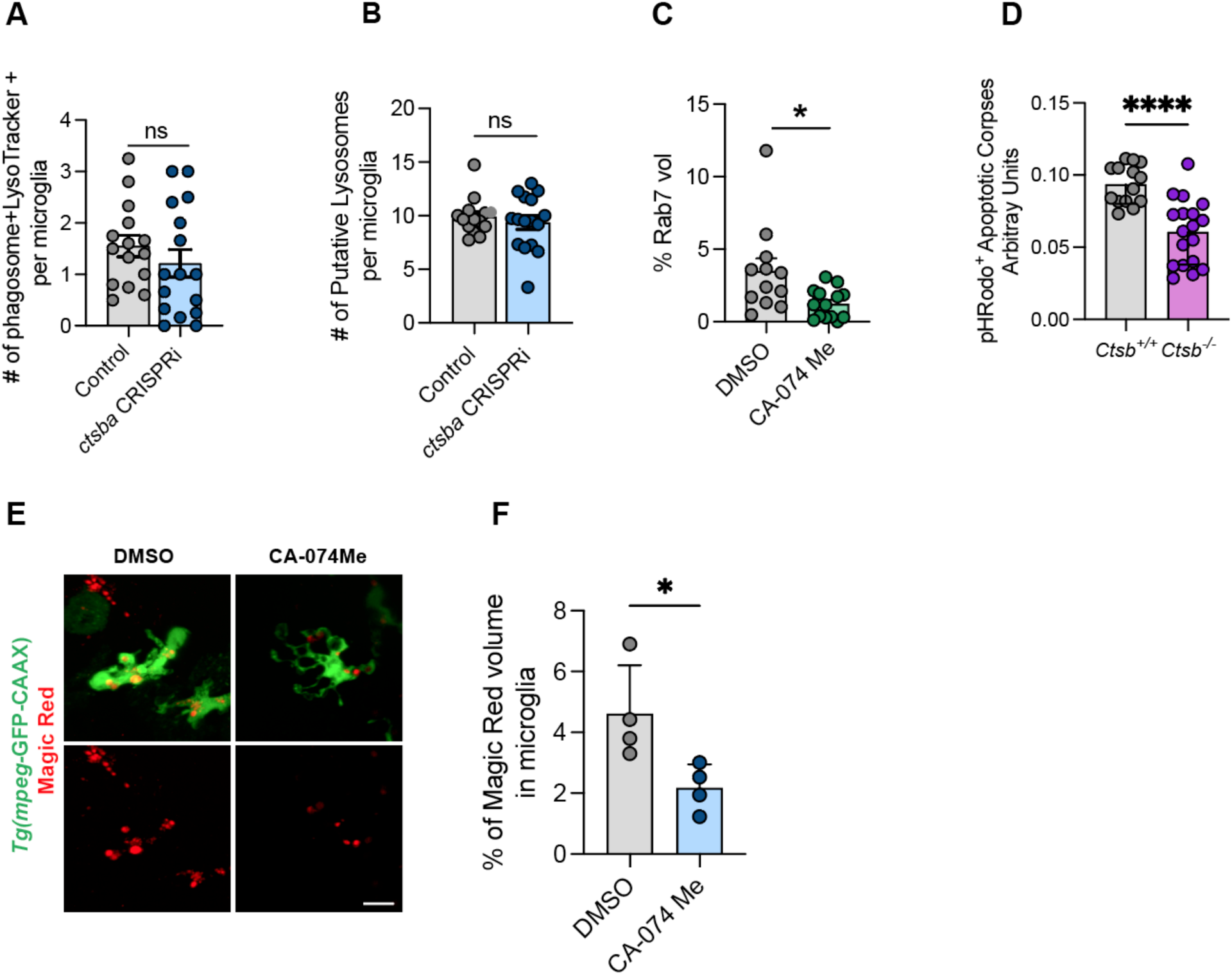
Characterization of microglial phagocytosis, related to Figure 3. **(A)** Number of acidified phagosomes (size > 3 µm) within microglia from wild-type and *ctsba* CRISPRi mutants. Welch’s t-test, ns. Dots =16 fish/group averaging 2-3 microglia per fish. **(B)** Number of putative lysosomes in control and *ctsba* CRISPRi mutants at 10 dpf. Size exclusion ≤ 2µm. Welch’s t test, ns. Dots =16 fish/group averaging 2-3 microglia per fish. **(C)** Total volume of Rab7-mCherry expression within *Tg(mpeg1:GFP-CAAX*) positive cells from wild-type and *ctsba* CRISPRi mutants in the optic tectum at 10 dpf (n=12). Welch’s t-test, *p < 0.0291. Dots = 10 fish/group averaging 2-3 microglia per fish. **(D)** Quantification of pHRodo^+^ signal in primary cultured microglia from *Ctsb ^+/+^* and *Ctsb ^−/−^* mice at 12 hours following the seeding of pHrodo-labeled apoptotic SH-S5Y5 cells. Dots represent 14-18 wells per genotype pooled from 3 independent experiments. Welch’s t-test. P < **** 0.0001. **(E)** Representative images of cathepsin B-cleaved Magic Red in microglia from DMSO and CA-074Me treated fish. Scale bar = 10 µm. **(F)** Quantification of the % Magic Red within microglia from wild-type and *ctsba* CRISPRi mutants. Welch’s t-test, * p < 0.0478. Dots = 4 fish/group averaging 3-5 microglia per fish.

**Figure S4:**
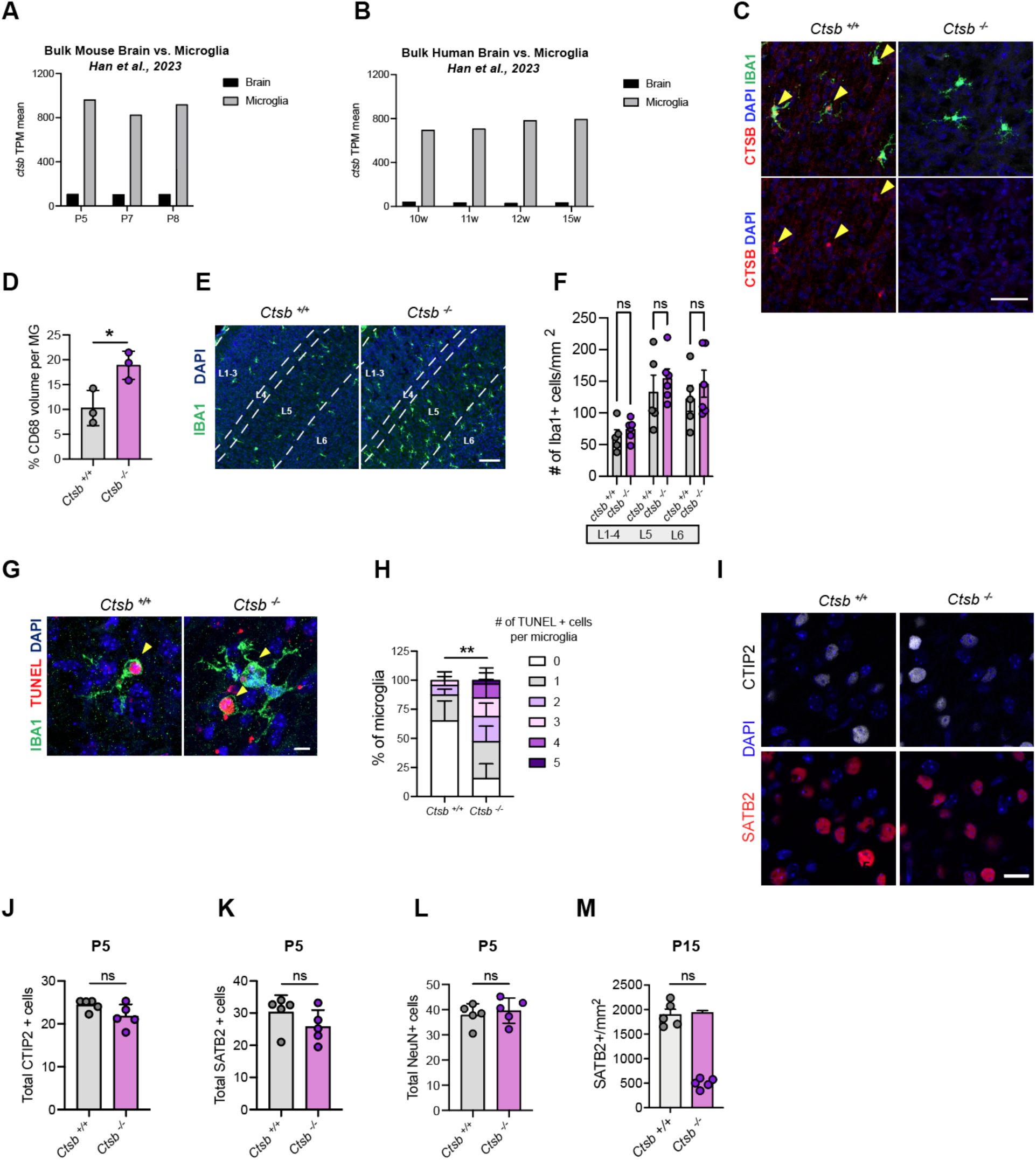
Additional characterization of *Ctsb*-deficient mice, related to Figure 4. **(A)** Expression of *Ctsb* from bulk sequencing of mouse brain vs. isolated mouse microglia at indicated postnatal ages (TPM: transcripts per million) from *Han et al., 2023.* **(B)** Expression of *CTSB* from bulk sequencing of human fetal brain and of microglia acutely isolated from human fetal brain at the indicate ages (w: gestational weeks) from *Han et al., 2023*. **(C)** Cathepsin B antibody in somatosensory cortex from *Ctsb ^+/+^*and *Ctsb ^−/−^* mice at P5 shows microglial-specific expression and antibody specificity. Scale bar = 25 µm. **(D)** Quantification of CD68 volume per microglia in the somatosensory cortex from *Ctsb ^+/+^* and *Ctsb ^−/−^* mice at P5. (n= 3 mice/group). Welch’s t-test, * p < 0.0327. **(E)** Representative images of IBA1^+^ staining in the somatosensory cortex from *Ctsb ^+/+^* and *Ctsb ^−/−^* mice at P5. (n= 5 mice for *Ctsb ^+/+^* and n= 6 mice for *Ctsb ^−/−^*). **(F)** Quantification of IBA1^+^ microglia cells in layers L1-L5 within the barrel cortex from *Ctsb ^+/+^* and *Ctsb ^−/−^* mice at P5. (n= 5 mice/group). 2-way RM ANOVA with Sidak’s multiple comparisons, ns. **(G)** Representative images of TUNEL positive cells within microglia from *Ctsb ^+/+^* and *Ctsb ^−/−^* mice. Yellow arrowheads indicate TUNEL inside microglia. Scale bar = 5 µm. **(H)** Quantification of TUNEL positive cells within microglia from *Ctsb ^+/+^* and *Ctsb ^−/−^* mice. Data calculated from means of 4-3 mice/group averaging 2-3 microglia per mouse. Fisher’s exact test, **p < 0.0066. **(I)** Representative images of CTIP2 and SATB2 neurons in the barrel cortex of P5 *Ctsb ^+/+^* and *Ctsb ^−/−^* mice at P5. Scale bar = 20 µm. **(J)** CTIP2^+^ neurons per mm^2^ in L5 in *Ctsb ^+/+^* and *Ctsb ^−/−^* mice at P5 (n=5 mice per group). Welch’s t-test, ns (0.107). **(K)** SATB2^+^ neurons density per mm^2^ in L5 in *Ctsb ^+/+^* and *Ctsb ^−/−^* mice, P5 (n=5 mice per group). Welch’s t-test, ns (0.205). **(L)** NeuN+ neuronal number in L5 in *Ctsb ^+/+^* and *Ctsb ^−/−^* mice at P5 (n=5 mice per group). Welch’s t-test, ns (0.5837). **(M)** SATB2^+^ neuron density per mm^2^ across all cortical layers in *Ctsb ^+/+^* and *Ctsb ^−/−^* mice at P15 (n=5 mice per group). Welch’s t-test, ns (0.7577).

## METHODS DETAILS

### EXPERIMENTAL MODEL AND SUBJECT DETAILS

#### Zebrafish

Wild-type, AB strain zebrafish (*Danio rerio*; ZIRC, University of Oregon, Eugene, OR) were propagated, maintained, and housed in recirculating habitats at 28.5°C and on a 14/10-h light/dark cycle. Embryos were collected after natural spawns, incubated at 28.5°C and staged by hours post fertilization (hpf). Larvae used were 10 days post fertilization (dpf), a time in development before sex determination. Ages were matched within experiments. The transgenic reporter lines, *Tg(mpeg1.1:EGFP-CAAX)*^50^ was used to mononuclear phagocytes and *Tg(mpeg1.1:Rab7-mCherry)*^21^ to label Rab7 expression in myeloid cells. All animal protocols were approved by and in accordance with the guidelines established by the Institutional Animal Care and Use Committee and Laboratory Animal Resource Center.

#### Mice

All mouse strains were maintained in the University of California San Francisco specific pathogen–free animal facility, and all animal protocols were approved by and in accordance with the guidelines established by the Institutional Animal Care and Use Committee and Laboratory Animal Resource Center. Mice were housed in a 12-hour light/dark cycle (7am-7pm) at 68-79°F and 30-70% humidity. Male and female mice were group-housed, when possible, with up to 5 mice per cage. Littermate controls were used for all experiments when feasible. The following mouse strain used is referenced in the text as *Ctsb^−/−^*: [B6;129-Ctsb^tm1Jde^*/J]* (Jax# 030971) mice^49^ were backcrossed six times to *C57BL/6J* and were obtained from Dr. Jason G. Cyster, University of California, San Francisco, CA. In all experiments, both male and female mice were used with ages specified in the legends.

### METHOD DETAILS

#### Magic Red injections

Larvae fish were injected with 2 nL of Magic Red at a dilution of (1:260) directly into the brain and returned to system water immediately following injections as previously described. Fish were imaged 4 hours post injection.

#### Molecular Biology

The CRISPRi plasmid was a gift provided by Dr. Cody Smith. The seed sequence for the gRNA targeting *ctsba* was the following: ACCATCTCATGGGACAAGGGAGG and identified using http://crispr-era.stanford.edu/ public software.

#### Generation of *ctsba* cell-type specific zebrafish mutant

The tol2 based mpeg:Gal4-VP plasmid was a gift from Dr. Sarah Kucenas and co-injected with the UAS:CRISPRi-U6:*ctsba* plasmid and transposase mRNA for integration. Casper embryos were injected at the 1 cell stage with 12.5 nl/µl of each respective plasmid and 25 ng/µl of transposase mRNA. F1 mutant hybrids were in-crossed to establish F2 homozygous generation. Screening was carried out by FACS and qPCR described below.

#### Fluorescence activated cell sorting (FACS)

For cell-type specific CRISPRi validation, 10 dpf *Tg(mpeg1.1:EGFP)* zebrafish brain were dissected (10 zebrafish were pooled per sample). Briefly, the brain(s) (regions) were mechanically dissociated in isolation medium (1x HBSS, 0.6% glucose, 15 mM HEPES, 1 mM EDTA pH 8.0) using a glass tissue homogenizer (VWR). Subsequently, the cell suspension was filtered through a 70 µm filter (Falcon) and pelleted at 300 g, 4°C for 10 minutes. The pellet was resuspended in 22% Percoll (GE Healthcare) and centrifuged at 900 g, 4°C for 20 minutes (acceleration set to 4 and deceleration set to 1). Afterwards, the myelin free pellet was resuspended in isolation medium that did not contain phenol red. Prior to sorting on a BD FACS Aria III, cell suspension was incubated with DAPI (Sigma). After sorting, cells were spun down at 500 g, 4°C for 10 min and the pellet was lysed with RLT+ (Qiagen). After sorting, cells were spun down at 500 g, 4°C for 10 min and resuspended in PBS + 0.05% BSA (Sigma).

#### Quantitative PCR

To extract RNA from cells isolated by FACS, freshly sorted cells were pelleted at 500 g for 10 minutes at 4° and then resuspended in RLT Plus buffer (Qiagen 1053393). Cells were vortexed and frozen for at least one day at −80° before being thawed on ice and processed for RNA using a RNeasy Plus Mini Kit (Qiagen). Purified mRNA was converted to cDNA with the High Capacity cDNA Reverse Transcription kit (Life Technologies) and amplified using either the Fast SYBR Green Master Mix (Thermo Fisher 43-856-12) and a 7900HT Fast Real-Time PCR System (Applied Biosystems).

#### Immunohistochemistry

For mouse brain tissue collection, mice were transcardially perfused with 5-10mL of sterile 1X PBS followed by 5-10mL of 4% paraformaldehyde. Tissues were fixed overnight at 4°C in 0.1M phosphate buffered 4% paraformaldehyde, cryoprotected with 20% sucrose, and embedded in optimal cutting temperature (OCT) medium (Sakura Finetek USA, Torrance, CA). Immunohistochemistry (IHC) was performed as previously described (Silva et al., 2020). Briefly, 16 to 25-μm-thick sections were collected and mounted onto glass slides or 40um thick floating sections were collected into 0.1M PBS. Fish brain sections were washed in phosphate buffer saline with 0.5% Triton-x (PBST) and incubated with 20% heat-inactivated normal goat serum in PBST for 2 hours (NSS; Sigma-Aldrich, Corp.). Mouse brain sections were blocked in staining buffer (5% normal goat serum, 0.4% Triton-X, 1X PBS) for 1 hour. Primary antibodies in staining buffer were applied overnight at 4°C. Sections were then washed with PBST and incubated in secondary antibodies for 2 hours at room temperature. For CTIP2 and SATB2 immunostaining, sodium citrate antigen retrieval was performed. Briefly, sections were immersed in fresh sodium citrate buffer (10 mM sodium citrate, pH 6.0) at 98°C for 10 minutes and cooled at room temperature for 20 minutes. IHC was performed as described above.

#### TUNEL staining

TUNEL staining was performed using In Situ Cell Death Detection Kit, TMR red (Sigma-Aldrich, CAT# 12156792910). Glass mounted sections were postfixed at 4C for 20 minutes prior to blocking. Following secondary antibody incubation, slides were permeabilized with PBS containing 1% sodium citrate/ 1% Triton-X-100 at 4C, rinsed with PBS, and then were incubated with TUNEL reaction cocktail per kit instructions for 1 hour at 37 C.

#### Confocal microscopy on fixed tissues

Images were acquired with a Zeiss LSM 800 laser scanning confocal microscope with 405, 477, 561, 650 nm laser lines. For mouse sections, 8-bit images were acquired with 5x scan speed at 1024×1024 resolution, 0.8 µm Z step size, and 2X line averaging. Laser power and gain were consistent within experiments.

#### Microglia engulfment assay

Images were acquired with an LSM 800 Confocal Microscope (Zeiss) using the same parameters as described above. Imaris software (Bitplane) was used to generate 3D surface rendering of microglia, which were then masked for TUNEL or active caspase 3 channels within those microglia. Masked channels were then 3D rendered to obtain volume data. TUNEL or active caspase 3 engulfment was calculated per cell as the volume of the respective signal divided by the volume of the microglia.

#### Microglia CD68 volume

Z-stacks were collected on an LSM 880 confocal microscope with AiryScan (Zeiss) on Superresolution mode and a 63x objective (NA 1.4). Laser power and gain were consistent across each image. AiryScan processing was performed in Zen software (Zeiss) at a setting of 6 (‘‘optimal’’ setting). Images were analyzed using Imaris software (Bitplane) by creating a 3D surface rendering of individual microglia, thresholded to ensure microglia processes were accurately reconstructed, and maintained consistent thereafter. Microglia rendering was used to mask and render the CD68 channel within each microglia. CD68 volume per microglia was then calculated as the total volume of masked CD68 volume within the masked GFP volume.

#### Neuronal subtype counts

Images of the somatosensory cortex from P15 mice were acquired using 20x magnification (NA 1.0) and 5×1 tiling to cover all cortical layers in a single optical section. NeuN staining was used to determine the center Z stack plane of the tissue section. All images were acquired in the same anatomical region of the somatosensory cortex, using the dorsal hippocampus as a reference point. Using FIJI image analysis software, ROIs were drawn for each cortical layer using DAPI staining. NeuN, CTIP2, and SATB2 analysis was automated followed by manual correction. Images were first segmented to binary images using Moments thresholding for CTIP2 and Huang thresholding for SATB2. Prior to quantification, the water-shedding function was used to segment touching cells. Particle analysis was used to quantify positive neurons using a size restriction ( size 20 μm-infinity). Neuronal counts were divided by the cortical layer area in mm^2^ to generate density per layer. 1-3 sections were imaged and quantified per mouse.

#### Zebrafish Live imaging

For live imaging, *Tg(mpeg:EGFP-CAAX)* zebrafish larvae incubated with LysoTracker (10mM) for 45 min then were anesthetized with 0.2 mg/ml of tricaine in embryo medium and mounted in 1.2% low-melting agarose gel on a glass bottom 35-mm dish (MatTek) and covered with embryo water containing 0.2 mg/ml tricaine. Time-lapse image was performed on a Nikon CSU-W1 spinning disk/high speed widefield microscope. We took time-lapse images from the optic tectum collecting 40-60 µm z-stacks (step size: 0.5 µm) at 4 min intervals for 1 hr and 30 mins. The images were processed by FIJI software. For analysis, we grouped the acidification events into three categories: 1) Phagosomes that were ‘Already’ acidified indicated by LysoTracker positivity 2) Phagosomes that became ‘Newly’ acidified.

#### Locomotion Behavior

Behavior studies on 10 dpf larvae were conducted in a 96-well plate using the automated locomotion detection device Danio Vision system and EthoVision XT 11.5 software (DanioVision, Noldus Information Technology). Larvae were placed into the DanioVision chamber to habituate for 30 mins before recording locomotion for 30 mins. At least 40 larvae per group (control and *ctsba* CRISPRi mutants) were used for a total of 3 independent experimental trials.

#### Primary microglia culture and engulfment assay

Mixed glia cultures were generated from P2 pups from *Ctsb ^+/+^*or *Ctsb ^−/−^* and grown in T75 flasks with DMEM (Gibco 11965126) supplemented with heat inactivated 10% FBS (Gibco 10437028) and 1% Pen/Strep (Gibco 15140122) at 37°C 5% CO_2_. Media was changed the next day and cultures were grown for another 10-12 days. Microglia were detached by hitting the flasks 10x against the bench and plated in a 96-well plate at a density of 20 000 cells/well in 100 µl final volume (4-6 replicate wells per condition). Cells were stained with Trypan blue (Invitrogen T10282) and counted using a Countess 3 Automated Cell Counter (Invitrogen). The next day, microglia were treated with 10,000 pHRodo^+^ apoptotic corpses (ratio 1:2) were added to the microglia 10-15 min before the first image acquisition (T=0h). Images were acquired using an Incucyte S3 Live Cell Analysis Instrument (Sartorius) at 20x, 4 images per well, every hour for 24h total in 555nm red channel (apoptotic corpses within lysosomes) and phase contrast (microglia). Images were then thresholded using the Incucyte Software for Live Cell analysis and the integrated intensity of the red channel (apoptotic corpses within lysosomes) normalized to microglia surface area (determined with phase contrast) was used for analysis. This value was multiplied by 10^3^ and is plotted as “pHrodo intensity (arbitrary units)” in **Figures 3L and S3D**.

#### Generation of apoptotic cells

The human neuroblastoma cell line, SH-S5Y5, was maintained in DMEM (Gibco 11965126) culture medium supplemented with 10% FBS and 1% Pen/Strep. To generate apoptotic corpses, cells were grown to 80% confluence and treated with 20 µM Navitoclax (Bcl-2 family protein inhibitor,^36^) for 24h. We confirmed that > 90% of the cells in the supernatant were dead cells (Trypan Blue^+^) (data not shown). Supernatant containing the dead cells was collected and centrifuged at 1000g/5min/RT. Cells were resuspended in PBS at concentration of 10^6^/mL. 1μl of 1 mg/ml pHrodo-SE (stock solution in DMSO, Thermo Fisher P36600) was added per 50 ml of cell suspension. Cells were incubated 30 min at RT before washing twice in PBS before resuspending them in DMEM supplemented with 10% FBS and 1% Pen/Strep. Apoptotic cells were used immediately for microglia engulfment assays.

#### Behavioral assay – Whisker nuisance test

The whisker nuisance test paradigm was adapted from^27^. Briefly, mixed gender P15 littermate controls and CTSB KO mice were used for a minimum of 3 experiments. Each mouse was habituated to an individual cage on 3 consecutive days from P12 to P14. Using the wooden end of a long q-tip, we then performed a sham stimulation, presenting the q-tip in front of the head of the mouse without touching it for a duration of 2 min followed by 3 consecutive 2 min trials gently stroking the whiskers on the right side of the face continuously, separated by 1 min intervals. All trials were recorded with a camera for later analysis. Using the video recording, five categories of behavioral responses to the whisker stimulation were scored as detailed in the table below. These parameters reflect increased avoidance, fear and aggression towards the probe. Interpretation: Normal behavioral responses to stimulation were assigned a zero value, whereas meaningful abnormal behavioral responses were assigned a value of 2. The maximum whisker nuisance score is 10. High scores (6–10) indicate abnormal responses to the stimulation, in which the mouse freezes, becomes agitated or is aggressive. Low scores (0–3) indicate normal responses, in which the mouse is either soothed, curious or indifferent to the stimulation.

| BEHAVIORS | SCORE |  |  |
| --- | --- | --- | --- |
|  | 0 | 1 | 2 |
| **Response to stick** | Interested or ignore | Avoiding, anxious | Attacks, bites |
| **Evasiveness** | No evasive behavior | Runs away <50% | Runs away, moves to protect whiskers |
| **Grooming** | Normal grooming | No grooming | Irritated scratching, rubbing, pulling |
| **Freezing** | Walks around, exhibits curiosity | Frozen < 50% time | Frozen, defensive, fearful >50% time |
| **Breathing** | Normal range |  | Hyperventilating, gasping |

### QUANTIFICATION AND STATISTICAL ANALYSIS

Graphpad Prism 8.3.0 was used for all histological quantification analyses. Statistical tests are described in text and figure legends.

### KEY RESOURCE TABLE

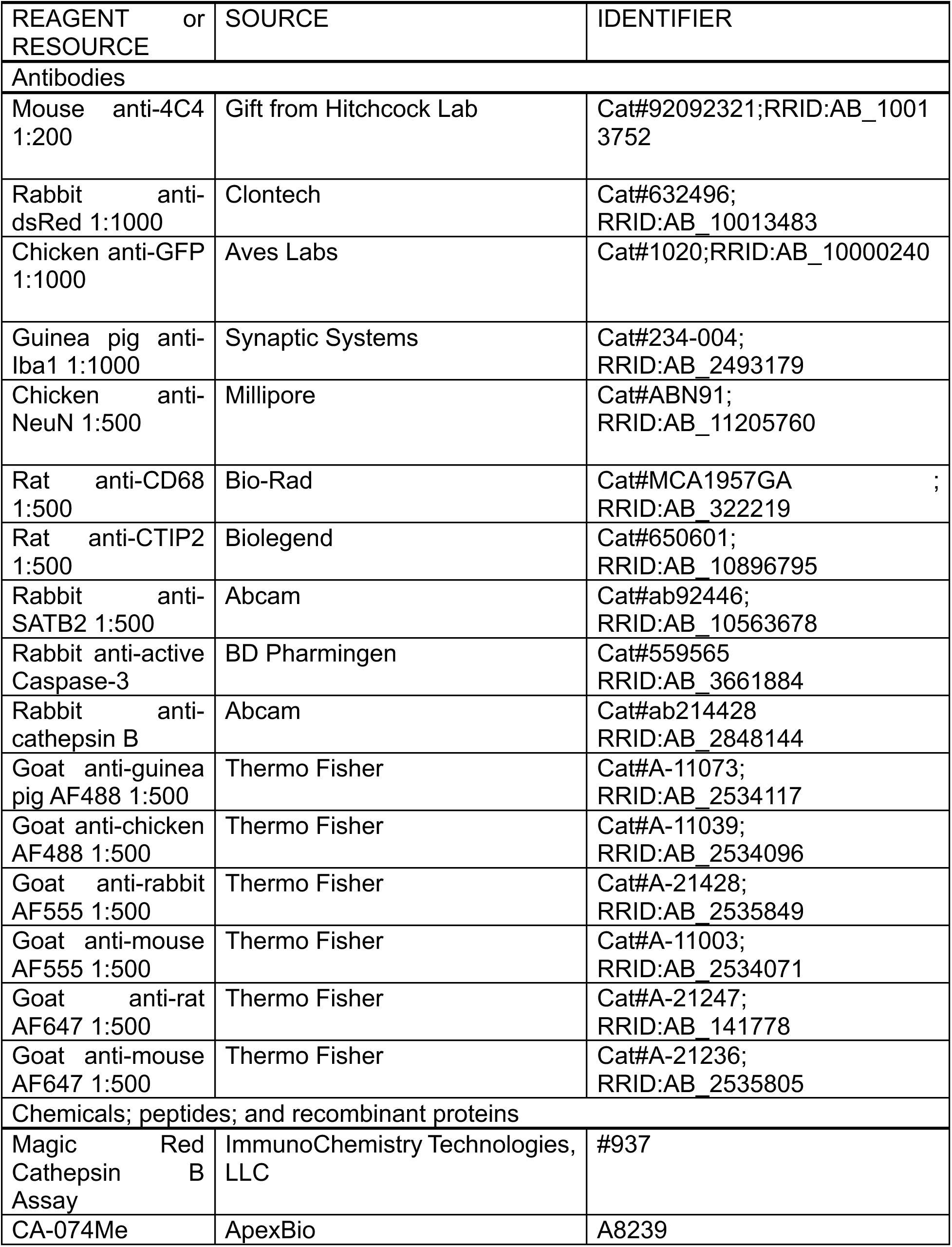
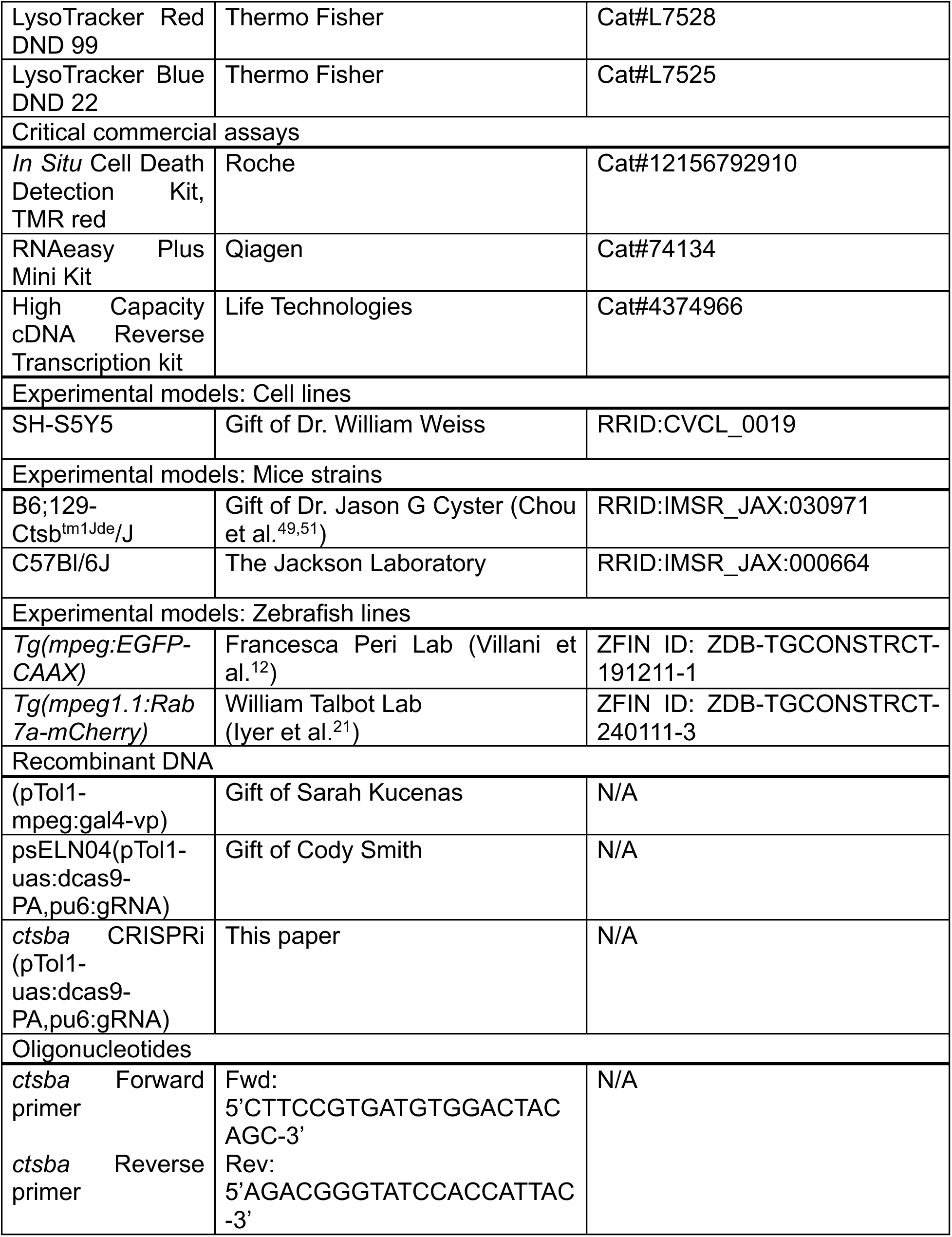
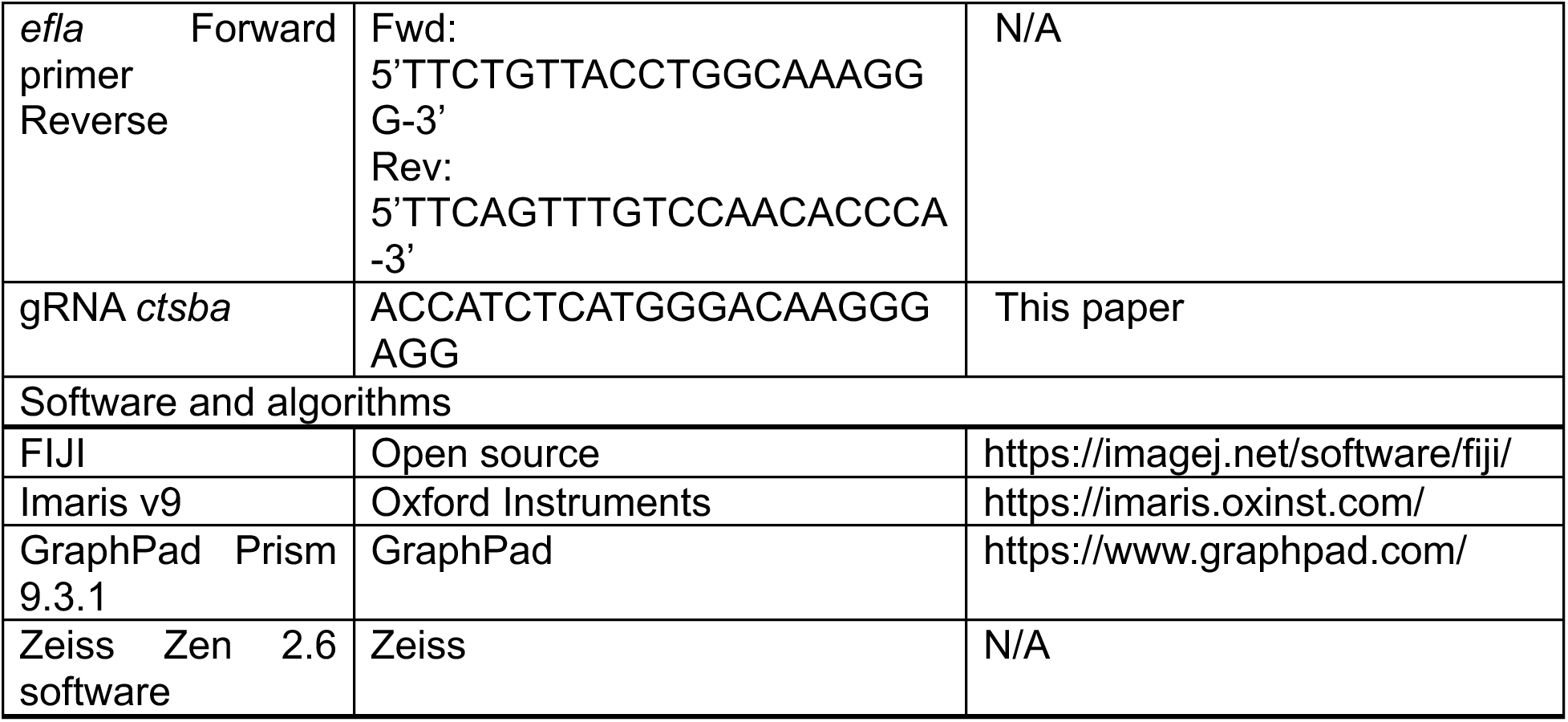

